# Evolution-Based Functional Decomposition of Proteins

**DOI:** 10.1101/022525

**Authors:** Olivier Rivoire, Kimberly A. Reynolds, Rama Ranganathan

## Abstract

The essential biological properties of proteins - folding, biochemical activities, and the capacity to adapt - arise from the global pattern of interactions between amino acid residues. The statistical coupling analysis (SCA) is an approach to defining this pattern that involves the study of amino acid coevolution in an ensemble of sequences comprising a protein family. This approach indicates a functional architecture within proteins in which the basic units are coupled networks of amino acids termed sectors. This evolution-based decomposition has potential for new understandings of the structural basis for protein function, but requires broad further testing by the scientific community. To facilitate this, we present here the principles and practice of the SCA and introduce new methods for sector analysis in a python-based software package. We show that the pattern of amino acid interactions within sectors is linked to the divergence of functional lineages in a multiple sequence alignment - a model for how sector properties might be differentially tuned in members of a protein family. This work provides new tools for understanding the structural basis for protein function and for generally testing the concept of sectors as the principal functional units of proteins.

## Introduction

The amino acid sequence of a protein reflects the selective constraints underlying its fitness and, more generally, the evolutionary history that led to its formation (1). A central problem is to decode this information from the sequence, and thus understand both the “architecture” of natural proteins, and the process by which they evolve. With the dramatic expansion of the sequence databases, a powerful strategy is to carry out statistical analyses of the evolutionary record of a protein family (2–6). With the assumption that the principal constraints underlying folding, function, and other aspects of fitness are conserved during evolution, the idea is to start with an ensemble of homologous sequences, make a multiple sequence alignment, and compute a matrix of correlations between sequence positions - the expected statistical signature of couplings between amino acids. Using mathematical analyses that explore different aspects of this matrix (7; 8), studies have exposed tertiary structural contacts in protein structures (Direct Coupling Analysis, or DCA, (4; 9)), determinants of binding specificity in paralogous protein complexes (5), and larger, collectively evolving functional networks of amino acids termed “protein sectors” (Statistical Coupling Analysis, or SCA (10). These different analytic schemes suggest a hierarchy of information contained in protein sequences that ranges from local constraints that come from direct contacts between amino acids in protein structures to global constraints that come from the cooperative action of many amino acids distributed through the protein structure. Sectors are interesting since they may represent the structural basis for functional properties such as signal transmission within (3; 6; 11–14) and between (15–17) proteins, allosteric regulation (6; 15; 18–20), the collective dynamics associated with catalytic reactions (16), and the capacity of proteins to adapt (21). In addition, experiments show that reconstituting sectors is sufficient to build artificial proteins that fold and function in a manner similar to their natural counterparts (22–24). Thus, the quantitative analysis of coevolution provides a powerful approach for generating new hypotheses about the physics and evolution of protein folding and function.

These results imply that together with structure determination and functional measurements, the evolution-based decomposition of proteins into sectors and local contacts should be a routine process in our study of proteins. However, the analysis of coevolution poses non-trivial challenges, both conceptually and technically. Conceptually, coevolution is the statistical consequence of the cooperative contribution of amino acid positions to organismal fitness, a property whose relationship to known structural or biochemical properties of proteins remains open for study. Indeed, there is no pre-existing model of physical couplings of amino acids with which to validate patterns of coevolution. Thus, the goal of coevolution based methods is to produce models for the pattern of constraints between amino acids that can then be experimentally tested for structural, biochemical, and evolutionary meaning. Technically, the analysis of coevolution is complicated by both the limited and biased sampling of sequences comprising a protein family. Thus, empirical correlations deduced from multiple sequence alignments do not always reflect coevolution. Interestingly, the complexities in sequence sampling can represent both sources of noise and useful signal in decomposing protein structures, and it is essential to understand these issues in effectively using methods of coevolution.

The DCA approach for mapping amino acid contacts has been well-described by analogy with established theory in statistical physics (25; 26). Here, we present the principles and implementation of the SCA method for identifying sectors and introduce new tools for understanding the global patterns of coevolution between amino acid positions. The methods are implemented in a python-based software package that is available to the scientific community, and illustrated in the main text using the small G protein family of nucleotide-dependent switches (27; 28) as a model system (see supplementary section V.A for complete tutorial). Two other examples are provided in supplementary sections V.B-C, with more analyses on our laboratory webpage. In prior work, we examined just the broadest level of coevolution to define sectors - quasi-independent groups of coevolving amino acids (10). We now go beyond this first-order decomposition to reveal a more elaborate internal architecture for sectors in which subgroups of amino acids diverge along functional, and sometimes phylogenetic, subfamilies within the sequence alignment. Overall, this work should provide the necessary basis for broad testing of the concept of protein sectors by the scientific community.

## Results

The SCA begins with an alignment (*M* sequences by *L* positions) representing a sampling of homologous sequences expected to share common selective pressures (Fig. 1 and supplementary section I.A). Standard sequence database searching algorithms (BLAST, PSI-BLAST, etc.) (29) together with automated alignment tools (e.g. PROMALS (30)), or available databases of multiple sequence alignments such as PFAM (31) seem to provide suitable sequence alignments. Since SCA concentrates on conserved features of protein sequences (see below), it is relatively robust to variations in alignment quality, but will depend on the depth and diversity of sampling of homologous sequences. While an alignment of a protein family is in principle sufficient for an analysis of coevolution, taxonomic and functional annotations and atomic structures are valuable for interpretation. In this work, we will assume that an atomic structure is known for at least one sequence in the alignment. We also assume that the alignment has been subject to a number of pre-processing steps in which positions and sequences with too many gaps are removed, and a simple sequence-weighting scheme is applied to correct for the trivial over-representation of high-identity sequences (see Box 1 and supplementary sections III.B and IV.A). With sequence weights, we can compute an effective number of sequences in the alignment *M′* = ∑_*s*_ *w*_*s*_ where *w*_*s*_ is the weight for sequence *s*. For computational efficiency, the alignment is then down-sampled to a limit that preserves *M′*. More advanced methods for treating sequence relationships are possible (32) and will require further study.

**FIG. 1.**
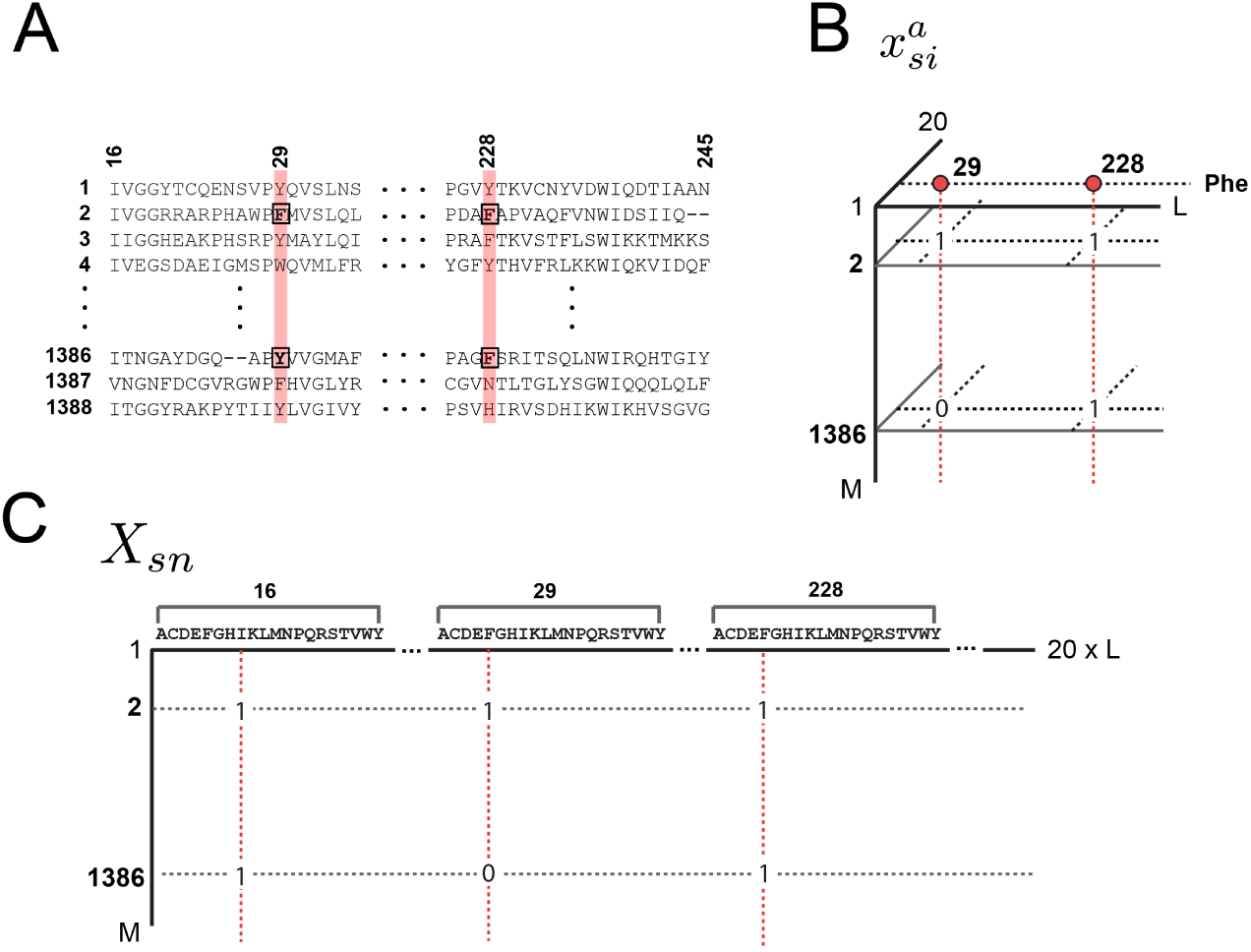
Three representations of a multiple sequence alignment comprised of *M* sequences and *L* positions. **A**, ascii text. **B**, a three-dimensional binary array 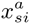, in which 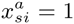 if sequence *s* has amino acid *a* at position *i*, and 0 otherwise; gaps are always set to 0. In this representation, the frequenci es of amino acids at individual positions are 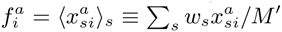, where *w*_*s*_ is the weight for each sequence *s* and *M′* = ∑_*s*_ *w*_*s*_ represents the effective number of sequences in the alignment. Joint frequencies of amino acids between pairs of positions are defined by 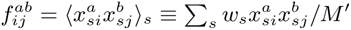. **C**, a two-dimensional alignment matrix *X*_*sn*_, in which the index *s* (along rows) represents sequences and the index *n* (along columns) represents the combination of amino acid and position dimensions in one, such that *n* = 20(*i* − 1) + *a*. This representation is useful in explaining the relationship between patterns of coevolution between amino acids and patterns of sequence divergence in the protein family (see Eq. (8)).

#### BOX 1: A SUMMARY OF CALCULATIONS

##### Alignment preprocessing

An alignment is represented by a *M × L ×* 20 binary array 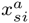 where *s* = 1, *…, M* labels the sequences, *i* = 1, *…, L* the positions, *a* = 1, *…,* 20 the amino acids, with 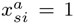 if sequence *s* has amino acid *a* at position *i* and 0 otherwise. Preprocessing steps:

1. Truncate excessively gapped positions based on a reference sequence or by a specified gap fraction cutoff (default, 0.4);
2. Remove sequences with a fraction of gaps greater than a specified value *γ* _*seq*_ (default, *γ* _*seq*_ = 0.2);
3. Remove sequences *r* with *S*_*r*_ < Δ, where *S*_*r*_ if the fractional identity between *r* and a specified reference sequence (default, Δ = 0.2);
4. Compute sequence weights *w*_*s*_ = 1/|{*r* : *S*_*rs*_ > *δ*}| w here *S*_*rs*_ if th e fractional identity between *r* and *s* (default, *δ* = 0.8), and truncate positions *i* with a frequency of gaps 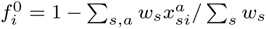 greater than a specified value *γ*_pos_ (default, *γ*_pos_ = 0.2); greater than a specified value *γ*_pos_ (default, *γ*_pos_ = 0.2);
5. Recompute the sequence weights *w*_*s*_ for the truncated alignment, and compute the frequencies of amino acid at individual positions *i* as 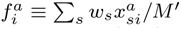, and at pairs of positions *ij* as 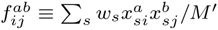, where *M′* = _*s*_ *w*_*s*_ represents the effective number of sequences in the alignment. When dealing with large alignments, a sixth step may added to speed up the subsequent calculations:
6. Resample *M″* sequences, with *M′ <M″< M*, by drawing them randomly from the original alignment with weights *w*_*s*_ so as to form an alignment with a smaller number of sequences but an equivalent effective number of sequences (which may slightly exceed *M*′, see SI; default, *M″* = 1.5 × *M*′).

##### Structure of evolutionary conservation

For a large and diverse alignment (*M′ >* 100, minimally), the evolutionary conservation of each amino acid *a* at position *i* taken independently of other positions is measured by the statistical quantity 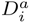, the Kullback-Leibler relative entropy of 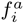 given *q*^*a*^, the background distribution of amino acids.

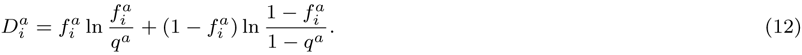

*q* is computed over the non-redundant database of protein sequences. If gaps are considered, and *θ* represents the fraction of gaps in the alignment, a back ground frequency for gaps can be taken as 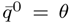, and then 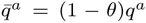 for the 20 amino acids. Also, 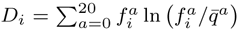 defines the overall conservation of position *i* taking all amino acids into account. To examine the co-evolution of pairs of amino acids, we introduce a measure that reports the significance of the raw correlations, 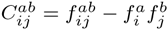, as judged by the degree of conservation of the underlying amino acids:

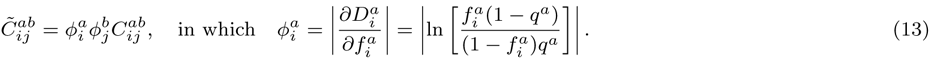

The information in the amino acid correlation matrix for each pair of positions is compressed into one number by computing the “Frobenius norm” of the 20 × 20 matrix 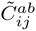 for each (*ij*):

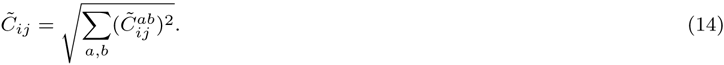

Analysis of 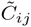 involves (a) spectral (or eigenvalue) decomposition of 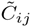, given by 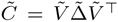, (b) determination of *k*\* significant eigenvalues (by comparison with vertically randomized alignments), (c) a transformation of the top *k*\* eigenvectors by independent components analysis (ICA), and (d) study of the pattern of residue contributions along independent components (ICs) 1 *… k\**. Distinct groups of positions can emerge along the ICs for two reasons: (1) the existence of multiple independent sectors, or (2) the hierarchicalbreakdown of one sector into subgroups that arise from global phylogenetic heterogeneities in the alignment.

##### Mapping and sector interpretation

The singular value decomposition (SVD) of the 20 × 20 matrix 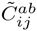 for each (*ij*), 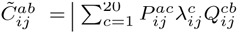, has the property that 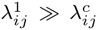 for *c* ≠ 1 (Fig. S1).That is, the information in the amino acid correlation matrix for each pair of positions can be compressed into one number, the top singular value (also known as the “spectral norm”). Besides compressibility, another empirical property of the SVD of 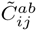 is that for a given position *i*, the top singular vector 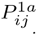 is (up to the sign) and therefore is essentially a property of just position *i* taken independently. This defines a projection matrix 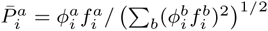 nearly independent of *j* (Fig. S4). That is, the amino acids by which a position *i* makes correlations with other positions *j* is nearly the same, with which we can reduce the *M × L ×* 20 alignment tensor 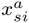 to an *M × L* alignment matrix 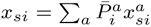. The matrix *x*_*si*_ gives a mapping between the space of positional correlations and the space of sequence correlations. Specifically, if 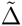 and 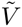 are the eigenvalues and eigenvectors of 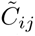, then

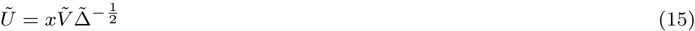

represents the structure of sequence space corresponding to the positional correlations in 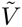. Also, if *W* is the transformation matrix derived from the ICA of 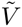, then 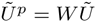 represents the sequence space corresponding to the ICs of 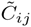. This mapping between position and sequence space provides a method to study the origin of the hierarchical pattern of coevolution that underlies sectors.

In this work, we use a PFAM-based alignment of the G protein family of nucleotide-dependent switches (PFAM, PF00071, version 27.0). The alignment comprises 21133 sequences by 1599 positions, but after the pre-processing steps with default values for thresholds, we obtain a final sequence alignment of 4978 sequences by 158 positions, with an effective number of sequences upon weighting of 3364. In what follows, we will assume that sequence weights are applied and for simplicity we will simply denote *M′* by *M*. No calculations below explicitly depend on its value; we shall only assume that *M′* is large enough to give good estimates of amino acid frequencies (*M′ >* 100).

An interesting point is that for nearly all alignments, the number of “variables” (*L*×20 possible amino acids) is typically on the order of or greater than the number of “samples” (*M*). Thus, it would seem impossible to reliably estimate the correlations between every pair of amino acids given such limited sampling. However, the sparsity of the constraints between amino acids observed both statistically (6; 10) and experimentally (33–35) effectively reduces the dimensionality of the solution, enabling practical approaches. The key issue is to propose a general approach for recognizing the “basis” -or groups of relevant amino acid positions -in which this solution largely exists. In SCA, the approach is to weight correlations by the degree of conservation of amino acids with the intuition that this fundamentally defines the relevance of features emerging from an evolutionary process. We develop this approach by first defining the first-order conservation of positions taken independently and then extending to correlations between positions.

### First-order statistics: position-specific conservation

The evolutionary conservation of a sequence position is estimated from the deviation of the observed distribution of amino acids at this position from a background distribution expected by neutral drift. A simple mathematical quantity that captures this concept is

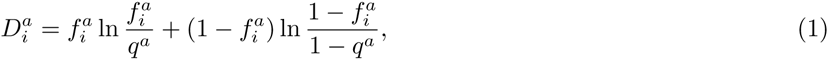

where 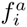 is the observed frequency of amino acid *a* at position *i* in the alignment and *q*^*a*^ is the background expectation (see supplementary section I.B for derivation). 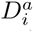 is known as the Kullback-Leibler relative entropy (36) and indicates how unlikely the observed frequency of amino acid *a* at position *i* would be if *a* occurred randomly with probability *q*^*a*^ – a quantitative measure of position-specific conservation. Note that 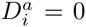 only when 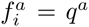 and increases more and more steeply as 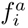 deviates from *q*^*a*^ (Fig. 2A), consistent with intuition that a measure of conservation should non-linearly describe the divergence of the observed distribution of amino acids from their expected values. An underlying assumption in the derivation of the relative entropy is that the sampling of sequences in the alignment is unbiased, a condition that, to varying extent, is violated by the tree-like phylogenetic structure of real alignments. But without validated models for protein evolution that can provide a basis for more accurate measures of conservation, this choice reflects the simplest definition that satisfies the general principle of conservation. Finally, Eq. (1) gives the conservation of each amino acid *a* at each position *i*, but an overall positional conservation *D*_*i*_ can be defined following the same principles (see supplementary section I.C).

**FIG. 2.**
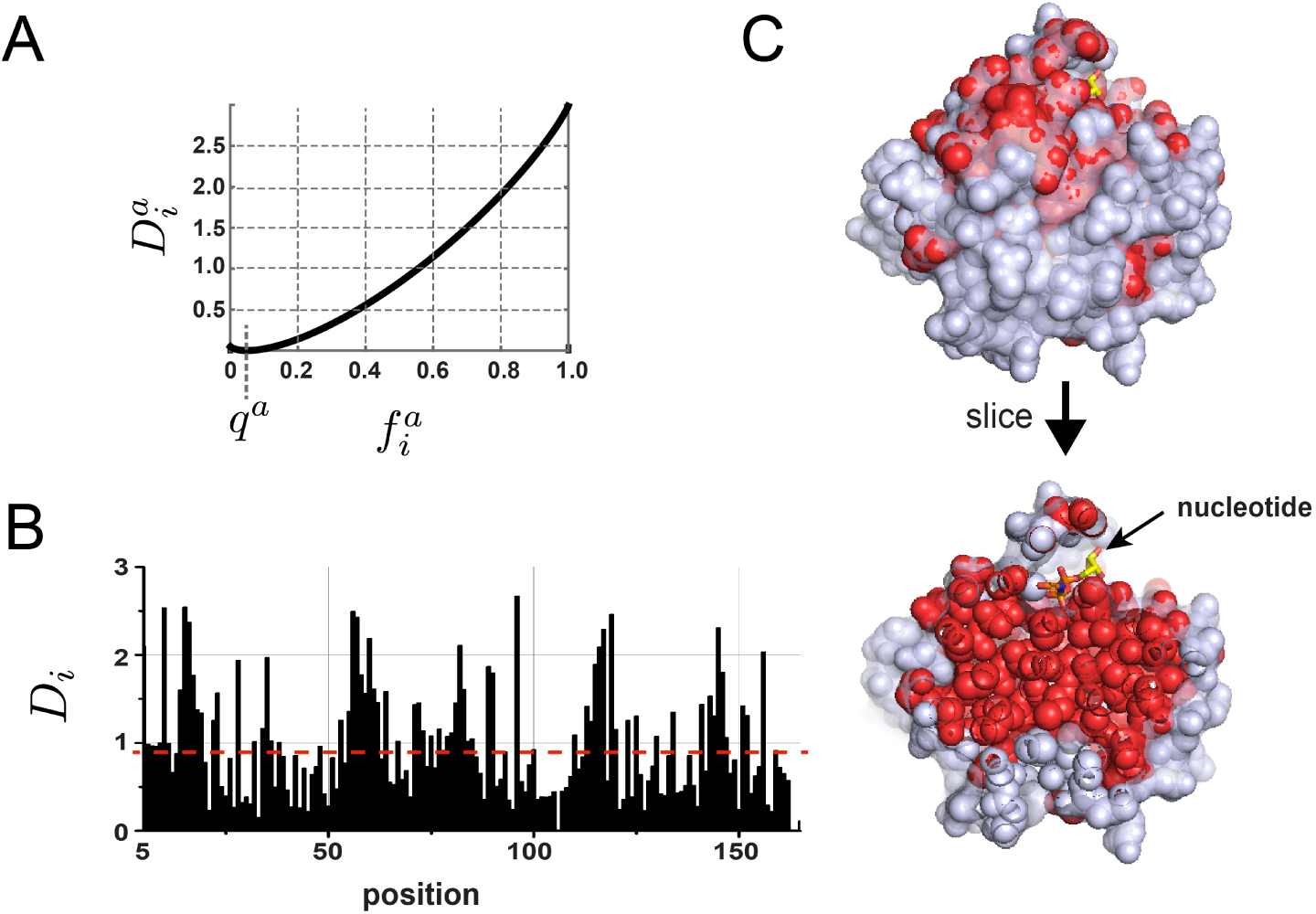
First-order conservation. **A**, A plot of 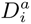, the measure of amino acid conservation, as a function of 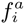, the amino acid frequency, and *q*^*a*^, the background frequency here for illustration set to 0.05. See the Supplementary Information for actual values of *q*. **B-C**, The overall positional conservation *D*_*i*_ for the G protein alignment, and a corresponding mapping on a slice through the core of the atomic structure of a representative member of the family (human Ras, PDB 5P21) showing that the top 50% of conserved positions (in red) lie at functional surfaces and within the solvent inaccessible core. Thus, positional conservation maps to a well-defined decomposition of protein structures: surface versus core.

Analysis of the spatial pattern of positional conservation generally leads to a simple conclusion: the solvent inaccessible core of proteins and functional surfaces tend to be more conserved and the remainder of the surface is less conserved (Fig. 2B-C) (10; 37; 38). Thus, positional conservation in sequence alignments reflects well-known properties of protein three-dimensional structures.

### Second-order statistics: conserved correlations

The cooperativity of amino acids in specifying protein folding and function implies that the concept of positional conservation of individual positions should at least be extended to a concept of pairwise conservation, reporting coevolution between positions in a protein family. Given the alignment, a measure of correlation of the pair of amino acids (*a, b*) at positions (*i, j*) is given by the difference of their joint frequency 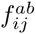 and that expected in absence of correlation, 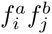. Computed for all pairs of amino acids in the alignment, this defines a covariance matrix

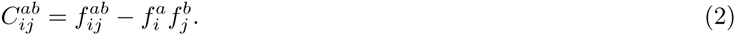

Alternatively, statistical dependency can be quantified by the mutual information, whose origin is similar to the relative entropy (5; 36). However, both the covariance matrix and the mutual information report deviations from independence given the frequencies 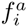, and do not take into account the evolutionary relevance of observing those frequencies. In the current implementation of SCA, the approach is to perform a first-order perturbation analysis on the multiple sequence alignment in which we compute the correlated conservation of pairs of amino acids. To explain, consider that many alignments *A* are available for the same protein family. We can then define relative entropies 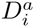,_*A*_-our measure of positional conservation - for each alignment *A*, and compute their correlations over the ensemble of alignments by

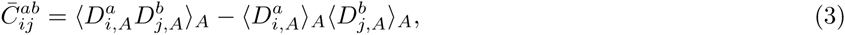

where the angled brackets indicate averages over the *A* alignments. In practice, many such alignments can be obtained by bootstrap resampling the original alignment (39); for instance, a procedure known as “jackknife resampling” consists of successively removing each sequence *s* from the original alignment to create a collection of *M* sub-alignments. Eq. (3) evaluated for these *M* “all-but-one” alignments yields a covariance matrix that has the form

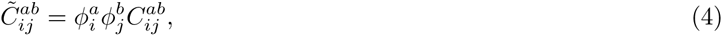

in which 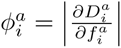 is a function of the conservation of each amino acid at each position (see supplementary sections I.D for derivation and IV.C for implementation) (10). The absolute value is taken to ensure positive weights. That is, SCA produces a weighted covariance matrix, with the weighting function *ϕ* controlling the degree of emphasis on conservation. This definition of *ϕ* has the property of rising steeply as the frequencies of amino acids 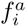 approach one. As a consequence, these weights damp correlations in 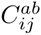 arising from weakly conserved amino acids (the gradient of 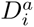 approaches zero as 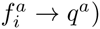, and emphasize conserved correlations.

Another way to understand these weights comes from considering their role in determining similarities between sequences comprising the alignment. The mathematical principles are described below, but in essence positional weights 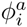 redefine the distance between sequences in a manner that emphasizes variation at more conserved positions in the alignment (see supplementary section I.I). It is logical that such a “conservation-biased” distance metric between sequences will provide a better representation of the functional differences (as opposed to historical differences) between sequences. The weighting by 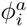 in Eq. (4) implements the same principle applied to the correlations between positions instead of the correlations between sequences.

In principle, the specific form of *ϕ* should vary depending on the evolutionary history of the protein properties that are under consideration; the more conserved the properties of interest are, the more the weights should emphasize conservation (40). Indeed, different weighting functions are possible if mathematical formalisms other than the KL entropy are proposed for defining positional conservation, or if other approaches than the first-order perturbation analysis described here are developed. These refinements might be justified with further experimental or theoretical understanding of the evolutionary divergence of protein function, but regardless, the result would be variation on the general theme in SCA of a weighted covariance matrix. Indeed, early versions of the SCA method (3; 6) involved slightly different weights whose technical origins are given in supplementary section I.E.

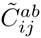 is a four-dimensional array of *L* positions × *L* positions × 20 amino acids × 20 amino acids, but we can compress it into a *L × L* matrix of positional correlations by taking a magnitude (the Frobenius norm) of each 20× 20 amino acid coevolution matrix for each pair of positions (*i, j*):

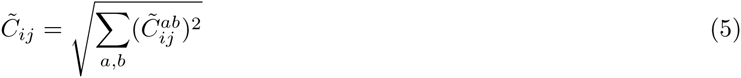

See supplementary section IV.C for implementation and section I.F and Fig. S1 for additional arguments about compressibility of 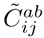.

Figure 3A-B shows the 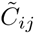 matrix for the G protein family. As previously reported, the matrix is heterogeneous, with a small number of positions engaged in significantly higher correlations than most positions (Fig. 3A, (6; 10)). Hierarchical clustering makes this heterogeneity more apparent (Fig. 3B). These findings are qualitatively consistent with a sparse, hierarchical, and cooperative pattern of evolutionary constraints. As shown below, in many cases there is also modularity (10), with quasi-independent clusters of positions emerging from the correlations (the sectors and their subdivisions). Unlike the conservation of positions taken independently (Figs. 2B-C), none of these properties is evident in current analyses of protein structures.

**FIG. 3.**
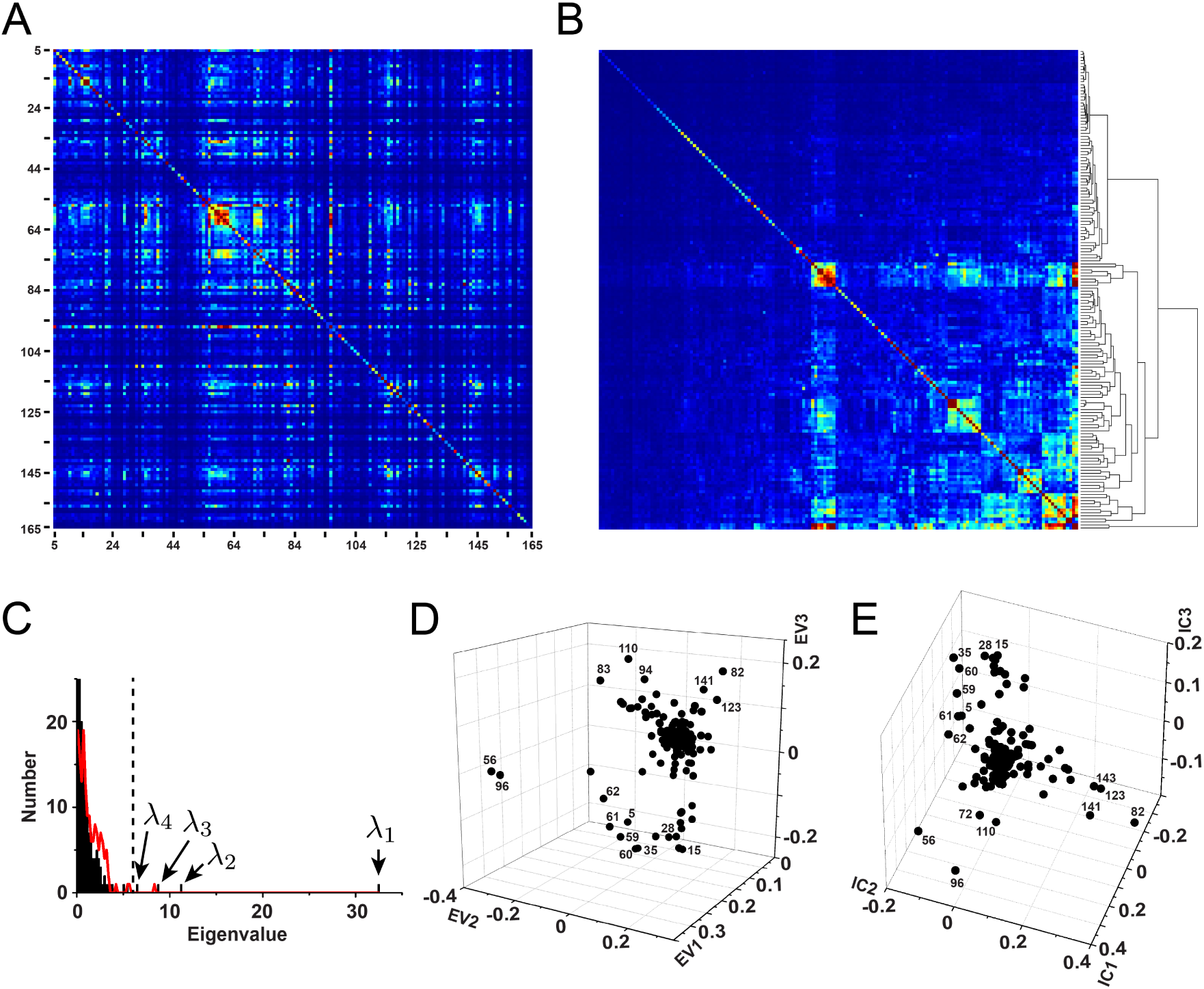
The SCA positional correlation matrix 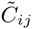 and spectral decomposition/ICA for the G protein family. **A-B**, 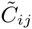 ordered by primary structure (**A**), and after hierarchical clustering (**B**). The data is consistent with a sparse and hierarchical organization of correlations – a general result for most protein families. **C**, The eigenspectrum of 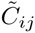 (in black bars), and the corresponding distribution expected randomly (in red) indicates that, conservatively, just the top four eigenmodes are relevant for further analysis. Note that the first random eigenvalue is ignored since it is a trivial consequence of retaining the independent conservation of sites in the randomization process (10). **D**, The top three eigenvectors suggest the possibility of distinct groups of coevolving positions, but illustrates the property that these groups emerge along combinations of eigenmodes. **E**, Independent components analysis (ICA) optimizes the independence of groups emerging along the different directions, putting the top three groups of amino acids on nearly orthogonal axes.

### Sector decomposition

How can we understand the pattern of correlations indicated by the 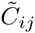 matrix (Fig. 3A)? An approach that is conceptually simple, computationally efficient, and well-suited for discovering low-dimensional patterns in high-dimensional datasets is spectral (or eigenvalue) decomposition. Per this decomposition, the 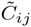 matrix is written as a product of three matrices:

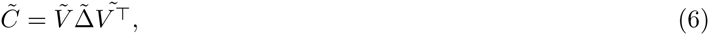

where 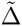 is an *L* × *L* diagonal matrix of eigenvalues (ranked by magnitude) and 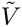 is an *L* × diagonal matrix of eigenvalues *L* matrix whose columns contain the associated eigenvectors. Each eigenvalue gives the quantity of information (variance) in 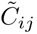 captured, and each associated eigenvector in 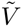 gives the weights for the contribution of sequence positions to the transformed variables (or eigenmodes). For the G protein alignment, the histogram of eigenvalues - the spectrum of 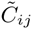 - reveals a few large eigenvalues extending from a majority of small values (Fig. 3C, black). To estimate the number of significant eigenvalues, we compare the actual spectrum with that for many trials of randomized alignments in which the amino acids at each position are scrambled independently (Fig. 3C, red line). This randomization removes true positional correlations, leaving behind the spurious correlations expected due to finite sampling in the alignment. As is the case for all practical alignments in which the number of effective sequences is not large compared to the number of amino acids, these spurious correlations generally account for the bulk of the spectrum. Indeed, in the G protein alignment this analysis indicates that just the top *k*\* = 4 eigenmodes are statistically significant. Thus, the *k*\* associated eigenvectors define a low dimensional space in which patterns of positional coevolution can be studied (Fig. 3D). It is important to note that the precise value of *k*\* is not a fundamental property of a protein family; it depends on protein size and the number of effective sequences. Nevertheless, with adequate sampling (*M′* > 100) the analysis of sectors seems largely robust to its precise value (see DHFR tutorial, supplementary section V.B).

The spectral decomposition is effective for dimension reduction, but the eigenmodes generally do not provide an optimal representation of groups of coevolving positions; instead, if independent groups are present, they typically emerge along combinations of the *k*\* top eigenvectors (Fig. 3D) (10). The reason is that just decorrelation of the positions by diagonalizing the 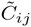 matrix - the essence of eigenvalue decomposition - is a weaker criterion than achieving statistical independence, which demands absence of not only pairwise correlations, but lack of any higher order statistical couplings. Independent components analysis (ICA (41; 42)) - an extension of spectral decomposition - is a heuristic method designed to address this problem. ICA uses numerical optimization to deduce a matrix *W* that transforms the *k*\* top eigenmodes of a correlation matrix into *k*\* maximally independent components (ICs, supplementary section I.G),

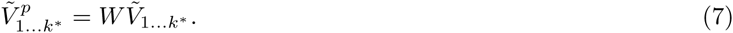

The bottom line is that the *k*\* ICs (in columns of 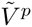) now represent a more appropriate organization of positional coevolution from which sector identifications can be carried out (Fig. 3E).

But, does the existence of *k*\* ICs imply *k*\* statistically independent groups (and therefore *k*\* sectors)? Not necessarily. Sectors typically have an organization in which the constituent positions can be further broken up into subsets of coevolving positions. As we show below, one generative mechanism for this architecture comes from the tree-like structure of the alignment in which sequences are partitioned into functional subfamilies along which portions of one sector can diverge (43; 44). Thus, when *k*\* is greater than the number of sectors, each IC could have one of two interpretations: (1) a truly independent sector associated with a distinct function, or (2) the decomposition of a single sector (representing one functional property) into sub-parts. In this sense, the term “independent component” is something of a misnomer, but we retain the language here for consistency with the ICA method.

How can we systematically distinguish these possibilities to deduce the number and composition of sectors? We follow a simple procedure (see supplementary section V, tutorials). First, we fit each IC to an empirical statistical distribution and identify the positions contributing to the top five percent of the corresponding cumulative density function (Fig. S2). The t-distribution appears to generally fit the ICs well in all cases studied to date. We then construct a sub-matrix of 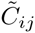 that contains only the selected top-scoring positions for the *k*\* ICs, ordered by their degree of contribution to each IC. For the G protein family, this corresponds to a matrix of 54 positions that comprises positions contributing to the four significant ICs (Fig. 4A). This sub-matrix describes both the pattern of “internal” correlations between positions that make up each IC (the diagonal blocks), and the pattern of “external” correlations between ICs (the off-diagonal blocks). Inspection of this ordered sub-matrix represents the basis for deducing the sector architecture. For the G protein family, IC4 shows near-independence from the other ICs, indicating that it defines a sector (sector 2, Fig. 4). In contrast, ICs1, 2, and 3 display strong inter-IC correlations, suggesting that these together comprise the hierarchically decomposed parts of a single sector (sector 1, Fig. 4). Fig. 4B shows a graphical representation of these conclusions, illustrating the hierarchical organization of all significantly coevolving positions into the *k*\* ICs.

**FIG. 4.**
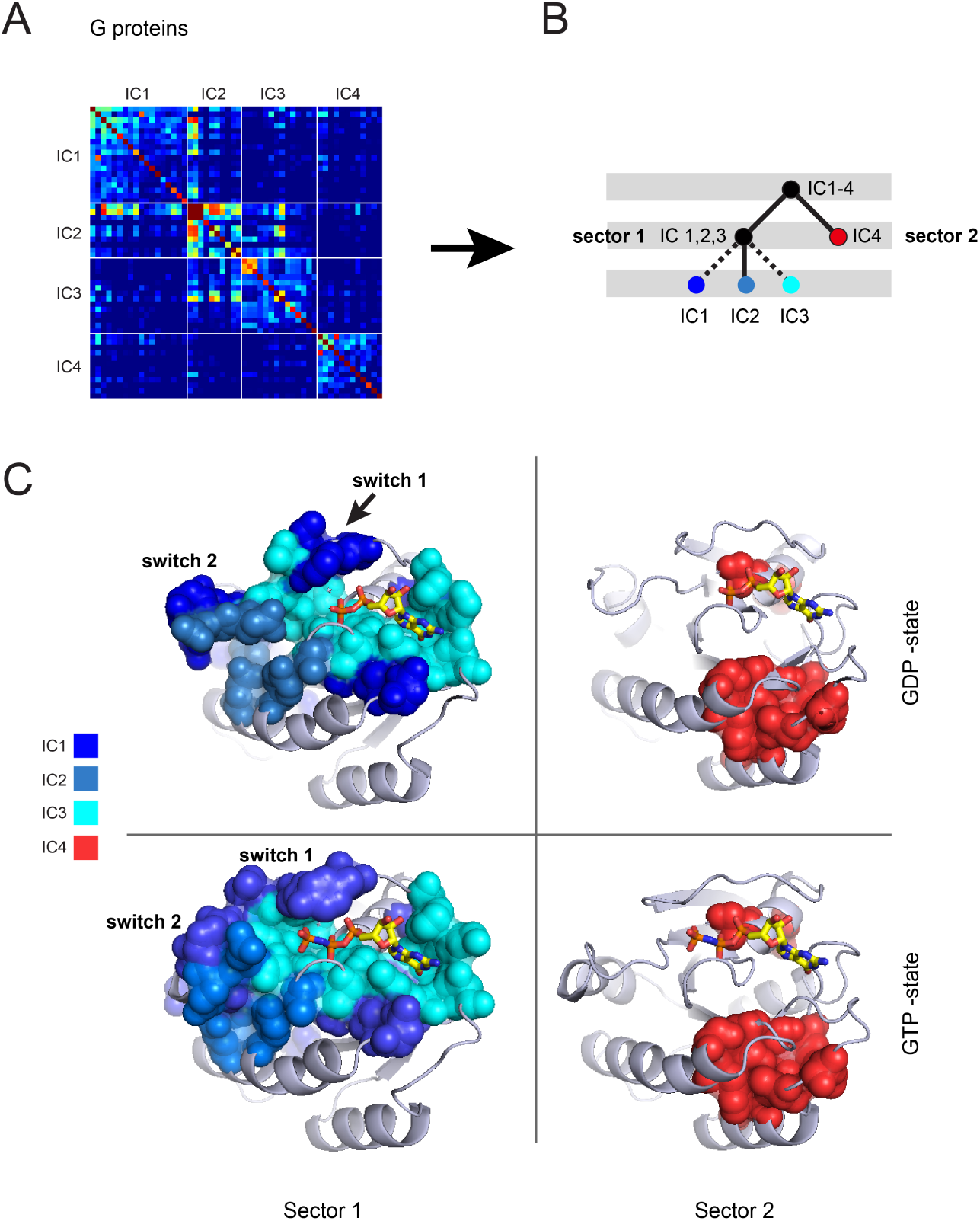
Sector identification for the G protein family. **A** shows the IC-based sub-matrix of the 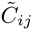 matrix and **B** shows a cartoon indicating the sector analyses. For the G protein family, IC4 represents a nearly independent group of coevolving positions (sector 2), while ICs 1, 2, and 3 show correlations between them that indicate the hierarchical parts of one sector (sector 1). **C**, Structural mapping of the G protein sectors. Sector 1 (blue colors) and sector 2 (red) mapped on atomic structures of the inactive GDP bound state of human Ras (PDB 4Q21), and the active GTP*γ*S bound state (PDB 5P21) as indicated. Overall, sector 1 corresponds well to the known allosteric signaling mechanism in the G protein family. The three quasi-independent ICs comprising sector 1 are represented in different colors as indicated, and are separately described in Fig. 5.

These sector definitions are made exclusively from the analysis of the IC-based sub-matrix of 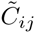, but mapping of the positions comprising the two sectors on representative atomic structures this protein family reveals an interesting spatial architecture (Fig. 4C). The guanine nucleotide-binding (G) proteins are a large family of binary switches that display different conformations depending on the identity of their bound nucleotide (27; 28). The essential characteristics of these proteins are exemplified by the high-resolution atomic structures of both inactive (GDP-bound, PDB 4Q21 (45)) and active (GTP*γ*S-bound, PDB 5P21 (46)) states of a canonical member of this family, H-Ras. The exchange of GTP for GDP triggers two specific conformational changes: clamping of the so-called switch I loop closer to the nucleotide binding pocket, and transit of a disordered and weakly interacting surface loop (switch II) to an ordered helix that is well-packed against the core domain (Fig. 4C) (27). Nucleotide exchange causes these conformational changes with little or no effect at other sites, indicating a specific and anisotropically distributed pattern of allosteric structural response limited to the switch domains.

Interestingly, mapping of the positions comprising sector 1 describes a physically contiguous group of amino acid residues that shows excellent agreement with the nucleotide-dependent allosteric mechanism (47). The sector is compact in the GTP-bound state but partially disrupted in the GDP-state, a finding consistent with the state-dependent connectivity between the nucleotide-binding pocket and the switch loops. Furthermore, the hierarchical breakdown of sector 1 into its constituent ICs 1, 2 and 3 reveals a meaningful structural decomposition: IC3 (cyan) defines a physically contiguous network that comprises the nucleotide binding pocket, IC1 (light blue) defines the packing interactions between switch II and the core domain, and IC2 (dark blue) represents a set of surface accessible positions (including switch I) that link to the buried core of sector 1. In addition, nucleotide exchange substantially reorganizes the structure and connectivity of IC1 and 2, but is largely inconsequential for IC3 (Figure 5). Consistent with assignment as an independent sector, sector 2 (red) also comprises a mostly physically contiguous group within the core of the G protein; like IC3 of sector 1 (cyan) it shows no nucleotide-dependent conformational plasticity. The physical role of sector 2 is unclear to our knowledge, but its definition represents a basis for further study. These findings show that in contrast to the first-order analysis of positional conservation, the study of conserved coevolution leads to new insights and hypotheses about protein function that are not merely a recapitulation of known structural or functional features. In the next section, we show that it also provides a link to the evolutionary history of the protein family. In addition to the G protein family (supplementary section V.A), further examples of sector analysis for two other protein families - the dihydrofolate reductases (48) and the class A beta-lactamases (49) - are provided in Fig. S3 and in tutorials (supplementary sections V.B-C).

**FIG. 5.**
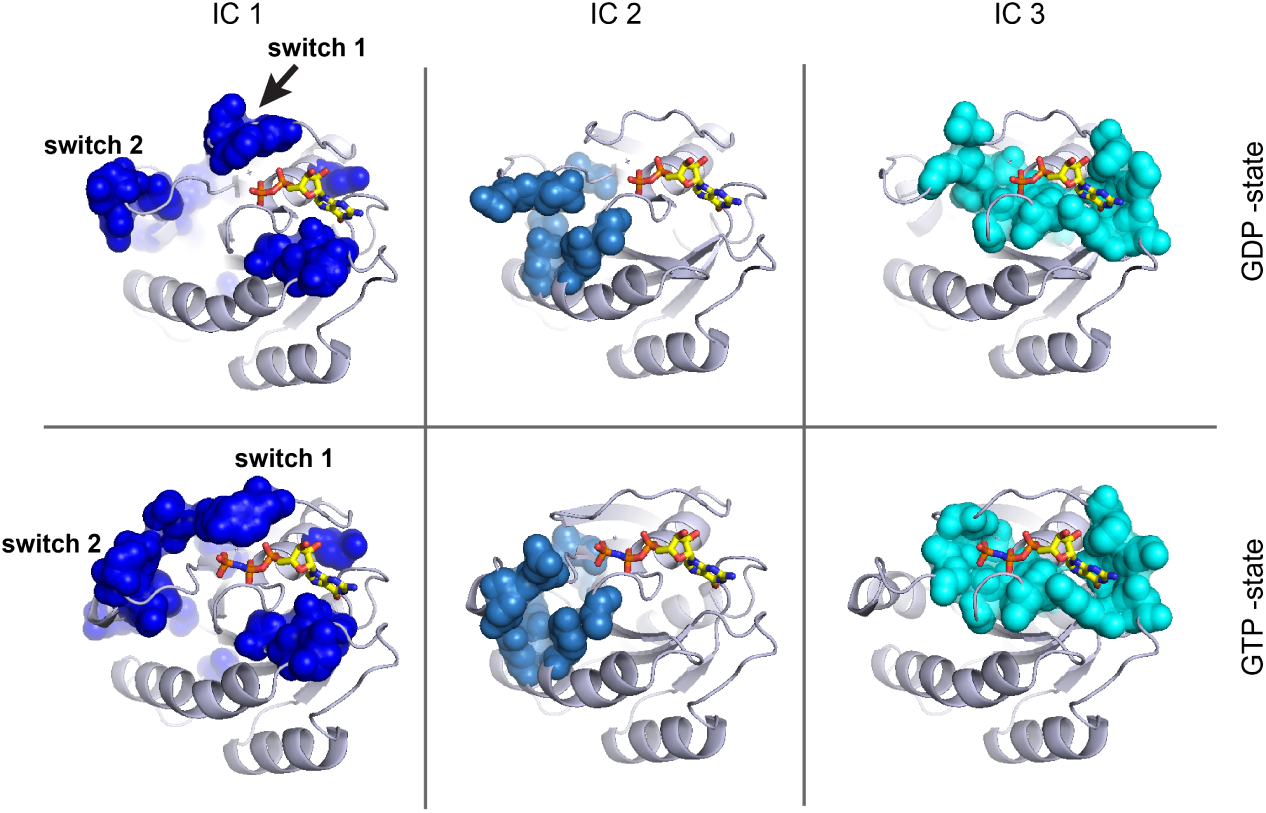
The state-dependent spatial organization of the three quasi-independent components comprising sector 1. The three ICs comprising sector 1 are shown on the atomic structure of the inactive GDP-bound state of human Ras (PDB 4Q21, top row), and the active GTP*γ*S bound state (PDB 5P21, bottom row). Like sector 2 (Fig. 4C), the IC3 component of sector 1 (cyan) shows little nucleotide-dependent conformational change, while ICs 1 and 2 of sector 1 show substantial conformational changes that are associated with allosteric switch regions I and II.

As presented here, the process of sector identification is made on examination of the IC-based submatrix of 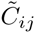 (Fig. 4A-B) and is heuristic, requiring the guided judgement of the practicing scientist to make the sector deductions. This choice is based on the need for broader experience with sector analysis to inform a more algorithmic and ultimately, automated approach for interpreting the 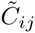 matrix. It also reflects the fact that groups of amino acids may coevolve at various degree and a continuity of situations may exist between two completely independent groups (two sectors) and a single one (a single sector). More generally, however, the current state of sector identification high-lights the limitations of spectral decomposition and ICA in inferring the hierarchical organization of the correlations.

However, given the general importance of this problem in many fields (50–52), it seems reasonable that the process of sector identification can be automated with further work.

### Sequence subfamilies and the basis of sector hierarchy

What underlies the hierarchical organization of protein sectors? A key insight comes from understanding a fundamental mathematical relationship between the pattern of positional correlations (defining sectors) and the structure of the sequence space spanned by the alignment (which defines sequence subfamilies) (43; 44). To explain, consider the two-dimensional binary matrix representation of an alignment *X*_*sn*_ comprised of *M* sequences by 20*L* amino acids (Figs. 1C and 6A). From *X*_*sn*_, we can compute two kinds of correlations: (1) a correlation matrix over rows 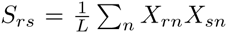, which represents the similarity (fraction identity) of each pair of sequences *r* and *s* (Fig. 6B) and (2) a correlation matrix over columns 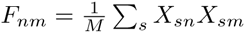, which represents the joint frequency of amino acids at each pair of positions (Fig. 6C). *F* and *S* are intimately related to each other by a mathematical property of the alignment matrix *X* known as the singular value decomposition (SVD). Specifically, if *U* represents the eigenvectors of the sequence correlation matrix *S* and *V* represents the eigenvectors of the amino acid correlation matrix *F*, then

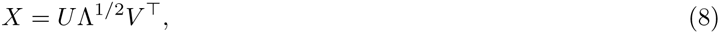

**FIG. 6.**
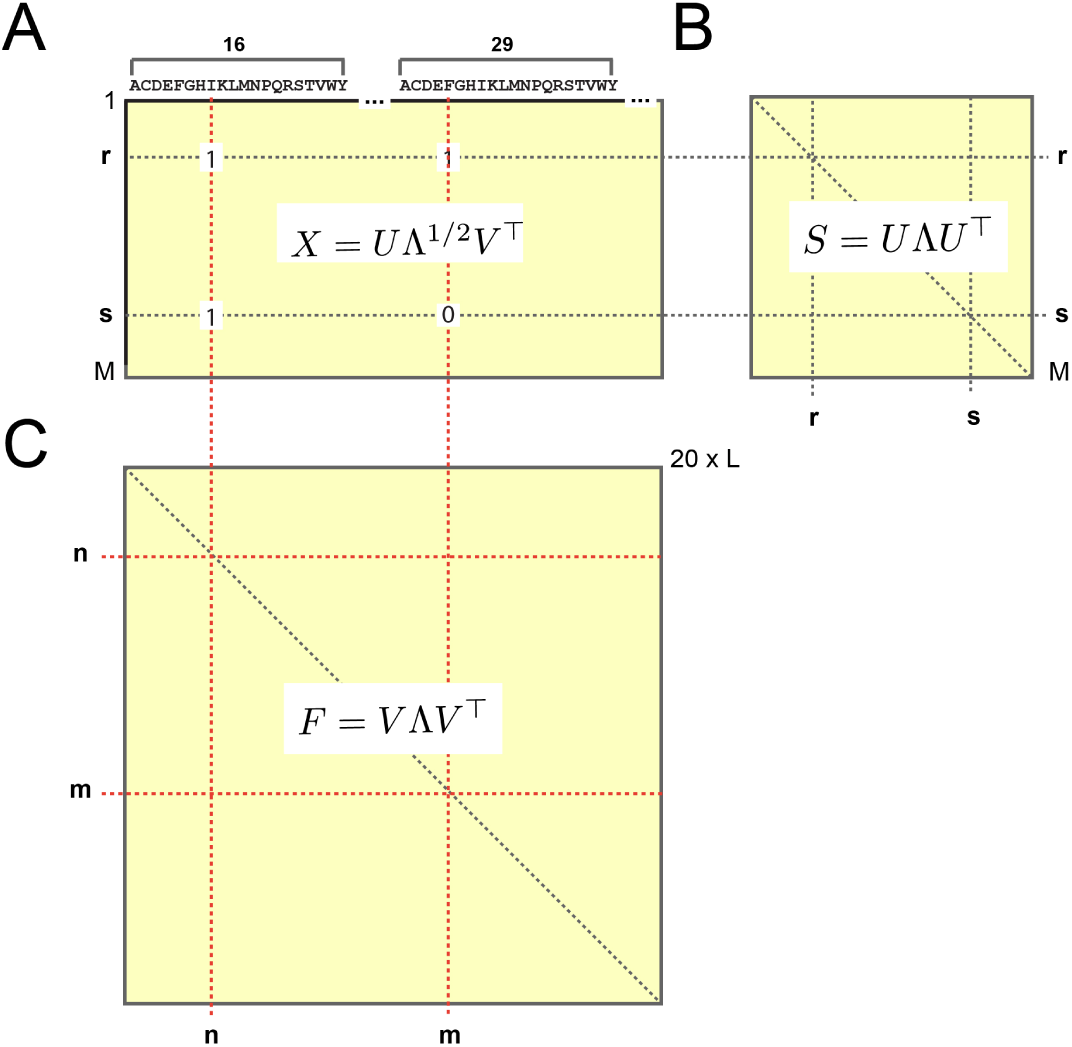
The mathematical relationship between sequence and positional correlations. **A**, A binary matrix representation of the alignment *X*_*sn*_, comprised of *M* sequences by 20× *L* amino acids (Fig. 1C); the equation shows the singular value decomposition (SVD) of *X* (Eq. (8)). From the alignment matrix, two correlation matrices can be computed: *S*, a correlation matrix over rows (**B**) describing relationships between sequences, and *F*, a correlation matrix over columns (**C**) describing relationships between amino acids; equations show the eigenvalue decompositions of these matrices. By the SVD, *X* provides a mapping between the two such that the eigenvectors of *F* (in *V*) correspond to the eigenvectors of *S* (in *U*). Thus, it is possible to associate coevolving groups of amino acids to patterns of sequence divergence in the alignment. As described in the text, a similar mapping is possible for positional (rather than amino acid specific) coevolution (Eq. (10)).

where Λ is a diagonal matrix whose entries are eigenvalues of both *S* and *F*. The key conceptual point is that by the SVD, the eigenvectors of *S* are a mapping from the eigenvectors of *F*, where the “map” is the alignment *X* itself,

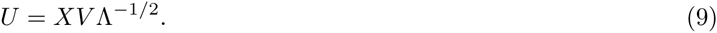

This introduces the principle of sequence/position mapping, using the full alignment matrix *X* to relate patterns of amino acid correlations (in *V*) to patterns of sequence divergence (in *U*). But, to study the pattern of sequence divergence associated with sectors, we need to make a similar mapping using the conservation-weighted dimension-reduced coevolution matrix 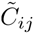 (rather than the unweighted amino acid correlation matrix *F*). Since 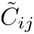 is a *L × L* positional correlation matrix, a sequence-space mapping analogous to Eq. (9) requires a dimension-reduced alignment matrix in which the 20 amino acids at each position are compressed into a single value. The Supplementary Information describes an approach for this step, effectively reducing the alignment 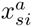 from a *M × L ×* 20 array to an *M × L* matrix *x*_*si*_ by projecting the amino acid dimension down to a single scalar value (supplementary section I.H and Fig. S4). By analogy with Eq. (9), the reduced alignment matrix *x*_*si*_ then defines a mapping between the space of positional coevolution (in the top ICs of the 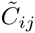 matrix) and the corresponding sequence space. Specifically, if 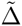 and 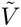 are the eigenvalues and eigenvectors, respectively, of the SCA positional coevolution matrix 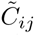, then

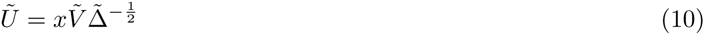

represents the structure of the sequence space corresponding to the patterns of positional coevolution in 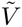. Furthermore, if *W* is the transformation matrix derived from ICA of 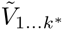, Eq. (7), then

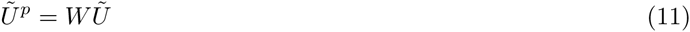

represents the sequence space corresponding to 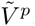, the ICs of the 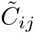 matrix.

The sequence/position mappings described in Eq. (10) and Eq. (11) give us the necessary tools to understand the origin of the hierarchical breakdown of sectors. Analysis of the G protein family revealed two sectors, one defined by a combination of contributions from ICs 1, 2, and 3, and one defined by positions contributing to IC4 (Figs. 4 and 5). Figure 7A-D shows the mapping between these four ICs and the corresponding sequence space, colored either by annotated functional sub-type of G protein (Fig. 7A-B) or by taxonomic origin (Fig. 7C-D). The data show that none of the ICs are associated with the divergence of the main taxonomic groups in the alignment; indeed, all taxa seem nearly homogeneously distributed over the sequence modes (*U ^p^*) corresponding to all the ICs.

**FIG. 7.**
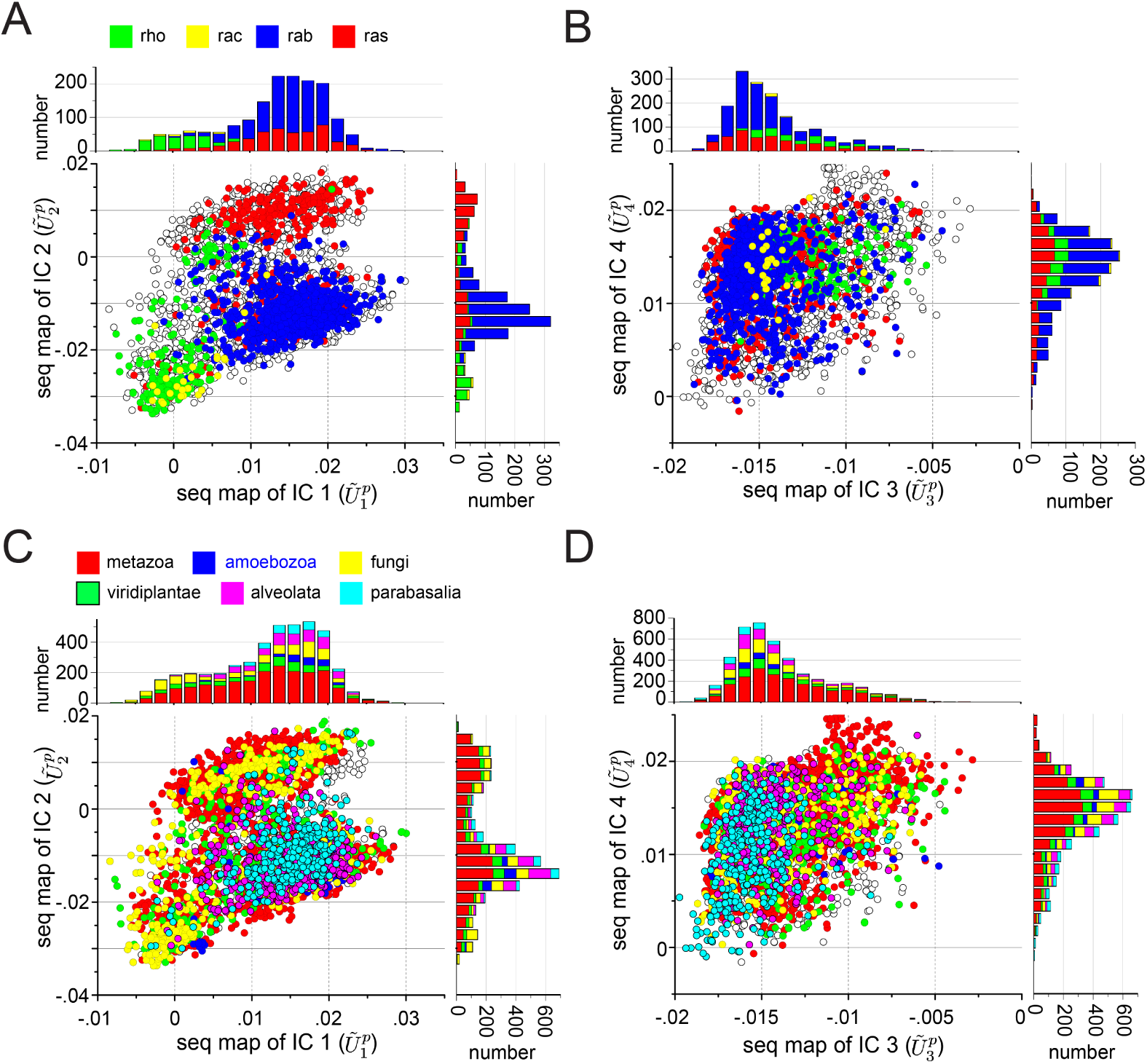
The relationship of positional coevolution to sequence divergences in the G protein family. The panels show scatterplots of sequences in the G protein alignment along dimensions(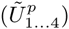) that correspond to sequence variation in positions contributing to each of the four ICs of the SCA coevolution matrix. The mapping between positional coevolution to sequence relationships is achieved using the reduced alignment matrix *x*, as per Eq. (10) and Eq. (11). Sequences are colored either by annotated functional sub-type of G protein (**A-B**) or by taxonomic origin (**C-D**), and the stacked histograms show the distributions of these classifications for each dimension. The data show that ICs 1 and 2 (**A**) correspond to distinct sequence divergences of functional subtypes of G protein; for example, IC1 separates the Rho proteins (green) along 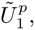, and IC2 separates the Rho proteins (green) and a subset of Ras proteins (red) along 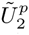. In contrast, IC3 and IC4 are homogenous with regard to G protein subtype (**B**), and all ICs are essentially homogeneous with regard to phylogenetic divergence (**C-D**). These data suggest that IC3 and IC4 are nearly homogeneous features of the G protein family, while IC1 and IC2 are differentially selected for more specialized properties of G protein subtypes.

What about classification by functional subtype of G protein? IC4 (sector2) and IC3 (cyan subset, sector 1) are also associated with a homogeneous distribution of functional subtypes (Fig. 7B) but IC1 (light blue subset, sector 1) and IC2 (dark blue subset, sector 1) separate different functional sub-classes (Fig. 7A). Specifically, IC1 separates the Rho-class of G proteins and IC 2 separates a subset of ras-like G proteins from the remainder. Thus, amino acid motifs within IC1 and IC2 have specifically diverged amongst different functional classes. This result is particularly interesting since IC1 and IC2 undergo significant nucleotide-dependent conformational change, make direct interactions with effector molecules, and are contiguous only in context of IC3. In contrast, IC3 and IC4 correspond to physically contiguous groups of amino acids that show no structural variation between the GTP-and GDP-bound states. These findings suggest the hypothesis that IC3 (cyan subset, sector 1) and IC4 (sector 2) are global functional modes shared by all members of the G protein family, while ICs 1 and 2 correspond to subsets of sector 1 that are specialized for tuning allosteric or effector-binding properties within sub-classes of G proteins. It will be interesting to test the idea that functional variations amongst G protein sub-types amounts to localized amino acid variations in small subsets of sector positions defined by the ICs.

It is interesting that none of the ICs in the G protein family show any heterogeneity with regard to the main taxonomic groups in the alignment (Fig. 7C-D). Many paralogs of the different functional classes of G proteins are found in each type of organism and thus the functional divergence of G proteins might not be expected to follow the divergence of species. In contrast, the breakdown of sectors is more associated with taxonomic classification for the DHFR protein family (Fig S2), consistent with the fact that this core metabolic enzyme is encoded by a single ortholog in each genome. See Fig. S5 and tutorials (supplementary sections V.B-C) for sequence/position mapping for the DHFR and beta-lactamase families.

Previous work has introduced the concept of sectors as quasi-independent units of protein structures that are associated with distinct functional properties (10), but has largely ignored their internal architecture. Here we present a more refined description in which a sector may itself be decomposed into a physically contiguous core element (e.g. IC3, Figs. 4C and 5), surrounded by peripheral elements (e.g. ICs 1 and 2, Figs. 4C and 5) that have the property of differential variation along functional branches of a protein family. These observations highlight the practical value of the mapping between units of positional coevolution and sequence subfamilies. When functional divergences between subfamilies are annotated, the mapping can identify the positions responsible for this divergence. For example, in the Hsp70 family of chaperones, the existence of subfamilies with known differences in allosteric function led to the identification of positions involved in the underlying mechanism (17). Turned around, when the role of specific positions in a protein is known, the mapping can help annotate sequences according to the associated functional property. For example, sequence divergence within sector positions with known function in the S1A family permitted classification of the sequence space according to that functional property (10). In principle, high-throughput methods for functional annotation of members of a protein family should permit even more refined mappings between amino-acid variation and phylogenetic or functional divergence, a step towards relating genotype-to-phenotype at the molecular level.

## Discussion

A fundamental goal in biology is to understand the architectural principles of proteins – the pattern of constraints on and between amino acids that underlies folding, biochemical activities, and adaptation. An emerging approach is to leverage the growing databases of protein sequences to statistically infer these constraints from large and diverse ensembles of homologous sequences. This strategy has two defining features that distinguish it from the more traditional direct physical study of specific model proteins. First, by averaging over the space of homologs, the statistical approach emphasizes the general constraints shared by many related proteins over those that are idiosyncratic to particular proteins. Second, by quantitatively examining the structure of correlations, the statistical approach provides models for the global pattern of cooperativity between amino acids. SCA adds an extra concept; by weighting correlations with a function of the evolutionary conservation of the underlying amino acids, this approach incorporates a measure of their functional relevance (53; 54). Mathematical decomposition of the weighted coevolution matrix reveals an internal architecture for proteins in which the basic functional units are groups of amino acids called sectors. The sector architecture is consistent with two empirically known but poorly understood properties of proteins: (i) *sparsity*, such that only a fraction of the amino acids are functionally critical (21; 55), and (ii) *distributed cooperativity*, such that folding and function can depend on the coupled action of amino acids linking distantly positioned sites (56–58). It has also revealed a previously unrecognized feature of proteins: *modularity*, such that multiple functionally distinct sectors are possible in a single protein domain (10). In this work, we extend this notion of modularity to show that sectors are themselves divided into quasi-independent subunits which are related to the divergence of selection pressures in functional subfamilies. These findings represent a basis for designing new experiments to understand the structural basis for protein function.

To the extent it is clear, it is valuable to explain the similarities and distinctions of SCA with other analyses of coevolution in multiple sequence alignments. The direct coupling analysis (DCA (4)) and its various extensions (9; 59– 61) are focused on using coevolution to determine physical contacts between amino acids within or between protein tertiary structures. As different as this problem may seem from discovering the pattern of *functionally* coevolving amino acids, there is a deep relationship. Recent work shows that the two approaches focus on two extremes of the same hierarchical architecture of coevolution (7; 8). SCA focuses on the global modes of coevolution (the top eigenmodes of a conservation-weighted correlation matrix), and DCA on the minimal units of coevolution (the bottom eigenmodes of an unweighted correlation matrix). Thus, coevolving direct contacts are at one end of the hierarchy and sectors at the other. Consistent with this, coevolving direct contacts are found within sectors and outside of sectors, but not bridging two independent sectors (8). Another approach for analyzing coevolution in protein alignments is mutual information, which has been successful at predicting the amino acids responsible for specificity in some protein-protein interactions (5). The distinction between this method and SCA lies in the nature of the weighting function *ϕ*; in essence, the mutual information method uses flat positional weights (*ϕ* = 1), which has the effect of emphasizing more unconserved correlations and may therefore be more appropriate when studying rapidly diverging functional properties (40). Taken together, these observations begin to clarify the relationship of the different approaches, and poses the question of the nature of physical information held at various levels of the hierarchy of coevolution, a matter for future experimentation. From a theoretical point of view, the observations highlight the need for a better, more unified framework representing the full hierarchy in amino acid correlations in proteins, a key next goal in advancing the statistical approach to the biology of proteins.

In general, sector analysis provides a representation of proteins that is distinct from the first-order analysis of positional conservation and that (so far) is not obtained from structure determination or functional mutagenesis. Thus, it provides a valuable tool for directing experimental studies of protein folding and function, and ultimately, for formulating a physical and evolutionary theory consistent with the design of natural proteins. Sector identification remains heuristic and a matter of scientific judgement for now, but our sense is that a rigorous algorithmic approach is matter of better definition of the problem itself, as experience grows with many case studies. The presentation of the SCA method and the associated software tools should facilitate the involvement of the scientific community in this process.

## Materials and Methods

Multiple sequence alignments were obtained from the PFAM database (release 27.0, accession codes PF00071 (G proteins), PF00186 (DHFR), and PF13354 (class A *β*-lactamases)), and were subject to pre-processing with default parameter values as described in 1. Reference sequences/structures selected for each family were human Ras (PDB 5P21), E. coli DHFR (PDB 1RX2), and E. coli TEM-1 *β*-lactamase (PDB 1FQG), and with sub-sampling to 2000 sequences, yielded the following final alignment statistics: G proteins (1809 effective sequences by 158 positions), DHFR (1178 effective sequences by 151 positions), *β*-lactamase (505 effective sequences by 200 positions). All calculations were carried out using a new python implementation of the statistical coupling analysis (pySCA v6.0), following the algorithms described in 1 and in the main text. Step-by-step tutorials for executing the analysis for the three protein families are provided in the Supplementary Information. The pySCA toolbox is available to the academic community for download through our laboratory website (systems.swmed.edu/rr_lab).

## Acknowledgments

We thank members of the Ranganathan Lab for critical review of the manuscript and I. Junier and S. Leibler for valuable discussions. This work was supported by Agence Nationale de la Recherche (ANR-10-PDOC-004-01, O.R.), the Gordon and Betty Moore Foundation (K. R.), the National Institutes of Health (RO1EY018720-05, R.R.), the Robert A. Welch Foundation (I-1366, R.R.), and the Green Center for Systems Biology.

